# MEK reduces cancer-specific PpIX accumulation through the RSK-ABCB1 and HIF-1α-FECH axes

**DOI:** 10.1101/2020.04.10.036103

**Authors:** Vipin Shankar Chelakkot, Kaiwen Liu, Ema Yoshioka, Shaykat Saha, Danyang Xu, Maria Licursi, Ann Dorward, Kensuke Hirasawa

**Author notes:** To whom correspondence should be addressed: Kensuke Hirasawa, Ph.D.; Division of BioMedical Sciences, Faculty of Medicine, Memorial University of Newfoundland, 300 Prince Philip Drive, St. John’s, NL, Canada, A1B 3V6.

## Abstract

The efficacy of aminolevulinic acid (5-ALA)-based photodynamic diagnosis (5-ALA-PDD) and photodynamic therapy (5-ALA-PDT) is dependent on the 5-ALA-induced cancer-specific accumulation of protoporphyrin IX (PpIX). We previously reported that inhibition of oncogenic Ras/MEK increases PpIX accumulation in cancer cells by reducing PpIX efflux through ATP-binding cassette sub-family B member 1 (ABCB1) as well as PpIX conversion to heme by ferrochelatase (FECH). Here, we sought to identify the downstream pathways of Ras/MEK involved in the regulation of PpIX accumulation via ABCB1 and FECH. First, we demonstrated that Ras/MEK activation reduced PpIX accumulation in RasV12-transformed NIH3T3 cells and HRAS transgenic mice. Knockdown of p90 ribosomal S6 kinases (RSK) 2, 3, or 4 increased PpIX accumulation in the RasV12-transformed NIH3T3 cells. Further, treatment with an RSK inhibitor reduced ABCB1 expression and increased PpIX accumulation. Moreover, HIF-1α expression was reduced when the RasV12-transformed NIH3T3 cells were treated with a MEK inhibitor, demonstrating that HIF-1α is a downstream element of MEK. HIF-1α inhibition decreased the activity of FECH and increased PpIX accumulation. Finally, we demonstrated the involvement of RSKs and HIF-1α in the regulation of PpIX accumulation in human cancer cell lines (DLD-1, SNB-75, Hs 578T, and MDA MB 231). These results demonstrate that the RSK-ABCB1 and HIF-1α-FECH axes are the downstream pathways of Ras/MEK involved in the regulation of PpIX accumulation.

## Introduction

The heme biosynthesis pathway, which is present in all cells, plays essential roles in cellular metabolism, including oxygen transport, regulation of cellular oxidation, and drug metabolism (1). Aminolevulinic acid (ALA) synthase is the first and a rate-limiting enzyme in the heme biosynthesis pathway. It synthesizes 5-aminolevulinic acid (5-ALA), which is then fluxed through the heme biosynthesis pathway, leading to the production of protoporphyrin IX (PpIX). PpIX is a fluorescent molecule and the immediate precursor of heme. As cancer cells generate high amounts of PpIX when treated with exogenous 5-ALA, PpIX fluorescence can be used for photodynamic diagnosis of tumors (5-ALA-PDD) (2–4). Moreover, irradiating PpIX with light of specific wavelength triggers the generation of reactive oxygen species (ROS), leading to cancer cell death. This is known as photodynamic therapy (5-ALA-PDT) (5). 5-ALA-PDD was recently approved by the US FDA for intraoperative optical imaging of tumors in patients with high-grade gliomas (6, 7), while 5-ALA-PDT has been in use in the clinic for treating non-melanoma skin cancer, non-small cell lung cancer, and cancers of the esophagus (8).

It is generally believed that oncogenic transformation promotes 5-ALA-induced PpIX accumulation in cancer cells. Transformation of mouse fibroblast cells with oncogenes such as K-ras or c-myc increased 5-ALA-induced PpIX accumulation (9). Similarly, Ras-transformed human mammary epithelial HB4a cells produced higher amounts of PpIX compared to parental HB4a cells when treated with 5-ALA (10). Furthermore, oncogenic transformation increases the expression of some enzymes in the heme biosynthesis pathway, including, porphobilinogen deaminase (PBGD), coproporphyrinogen-III oxidase (CPOX), and porphobilinogen synthase (PBGS), accelerating the synthesis of PpIX (11–13). These findings demonstrate that oncogenic transformation is a driver for PpIX accumulation.

In contrast, we identified that the activation of MEK, a downstream element of Ras, reduces 5-ALA-induced PpIX accumulation through two independent pathways – increased expression of ATP-binding cassette sub-family B member 1 (ABCB1), one of the PpIX efflux pumps, and increased activity of ferrochelatase (FECH), the enzyme that catalyzes the conversion of PpIX to heme (Fig. 1) (14). Furthermore, treatment with a MEK inhibitor significantly increased the efficacy of tumor diagnosis (5-ALA-PDD) and treatment (5-ALA-PDT) in animal models (14, 17). Importantly, the promotion of PpIX accumulation by MEK inhibition was cancer-specific, as it was not observed in normal cells *in vitro* and healthy organs *in vivo*. Our studies demonstrated that oncogenic activation of Ras/MEK is an excellent therapeutic target to promote the efficacy of 5-ALA-PDD and PDT. These findings are consistent with other studies that showed that MEK inhibition increases cancer cell death induced by 5-ALA-PDT (15) and that the activation of ERK, a downstream element of Ras/MEK, underlies cancer cell resistance to 5-ALA-PDT (16). Overall, Ras/MEK activation reduces PpIX accumulation leading to lower efficacy of 5-ALA-PDD and PDT. Considering that oncogenic transformation is known to accelerate PpIX accumulation, other oncogenic pathways or Ras downstream elements other than MEK may activate the enzymatic steps in the heme biosynthesis pathway leading to increased PpIX production. Nevertheless, further studies are required to understand the detailed molecular mechanisms by which oncogenic transformation regulates cellular PpIX accumulation.

**Fig 1.**
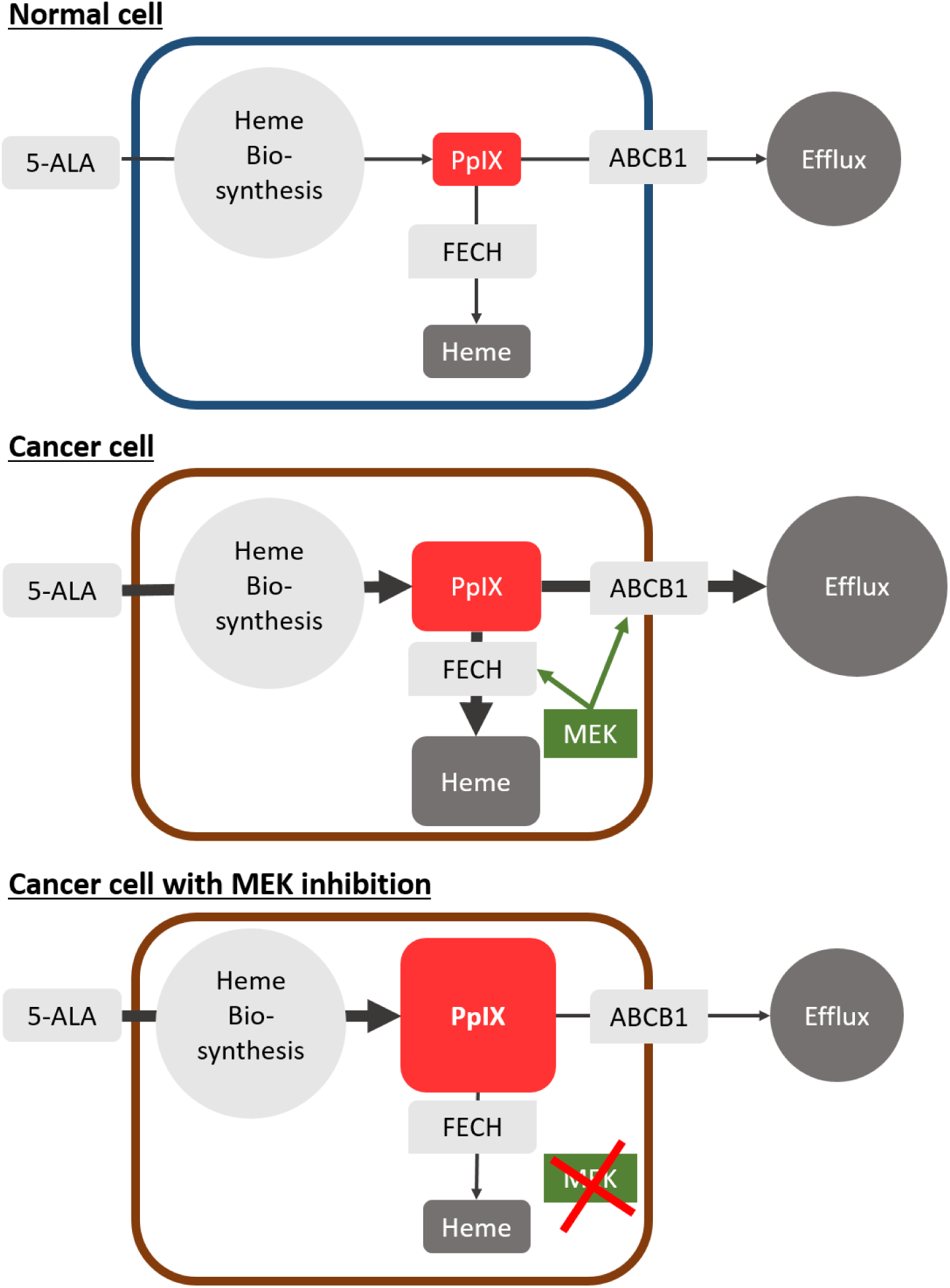
Schematic representation of protoporphyrin IX (PpIX) generation in normal and cancer cells. Upon exogenous stimulation of 5-ALA, cells generate PpIX via the heme biosynthesis pathway. PpIX is subsequently converted to heme by FECH or transported outside the cells through efflux receptors such as ABCB1. As oncogenic transformation activates enzymes of the heme biosynthesis pathway, cancer cells generate PpIX more efficiently than normal cells. As the Ras/MEK pathway promotes PpIX conversion to heme and PpIX efflux through ABCB1, MEK inhibition further enhances PpIX accumulation in cancer cells.

Activated Ras/MEK phosphorylates ERK 1 and 2, which in turn can activate multiple downstream signaling molecules such as MAP kinase-interacting kinases (MNKs), mitogen- and stress-activated protein kinases (MSKs), and 90 kDa ribosomal S6 kinases (RSK) (18). These downstream elements regulate cell proliferation, differentiation, and survival by modulating the transcription and translation of specific proteins. Although we previously demonstrated the involvement of MEK and ERKs in PpIX regulation (14), the Ras/MEK downstream elements responsible for the regulation of ABCB1 and FECH remains unidentified. Identifying the downstream elements is critical in gaining a better understanding of the cancer-specific accumulation of PpIX, the central cellular mechanism for 5-ALA-PDD and PDT. In addition, the identified downstream elements could be better therapeutic targets than upstream MEK for improving the efficacy of 5-ALA-PDD and PDT as they may have fewer off-target effects. Finally, they can be developed as prognostic markers to predict the efficacy of 5-ALA-PDD and PDT in clinical settings. To this end, in this study, we sought to identify the downstream elements of Ras/MEK that regulate PpIX efflux through ABCB1 and PpIX conversion to heme by FECH.

## Results

### Ras/MEK regulates 5-ALA-induced PpIX accumulation in mouse fibroblasts and transgenic mouse systems

In our previous studies, we used human cancer cell lines to demonstrate the regulation of PpIX accumulation via the Ras/MEK pathway (14, 17). As most human cancer cell lines have mutations that activate multiple oncogenic signaling pathways, it is often difficult to interpret the specific roles of a single activated pathway. Therefore, to eliminate the effects of other oncogenic signaling pathways on PpIX accumulation, we used RasV12-transformed mouse fibroblast NIH3T3 cell line, in which the Ras pathway is the only activated oncogenic pathway. As a first step, we confirmed that oncogenic activation of Ras/MEK reduces PpIX accumulation in the mouse fibroblast system. RasV12 cells were treated with or without three different MEK inhibitors (U0126, Selumetinib, or Trametinib) for 20 h and then with 5-ALA for 4 h (Fig. 2). Western blot analysis demonstrated that the amount of phosphorylated ERK was reduced in RasV12 cells treated with the inhibitors, suggesting that the Ras/MEK pathway was effectively inhibited. Treatment with all three MEK inhibitors significantly increased 5-ALA-induced PpIX accumulation in RasV12 cells in a dose-dependent manner.

**Fig 2.**
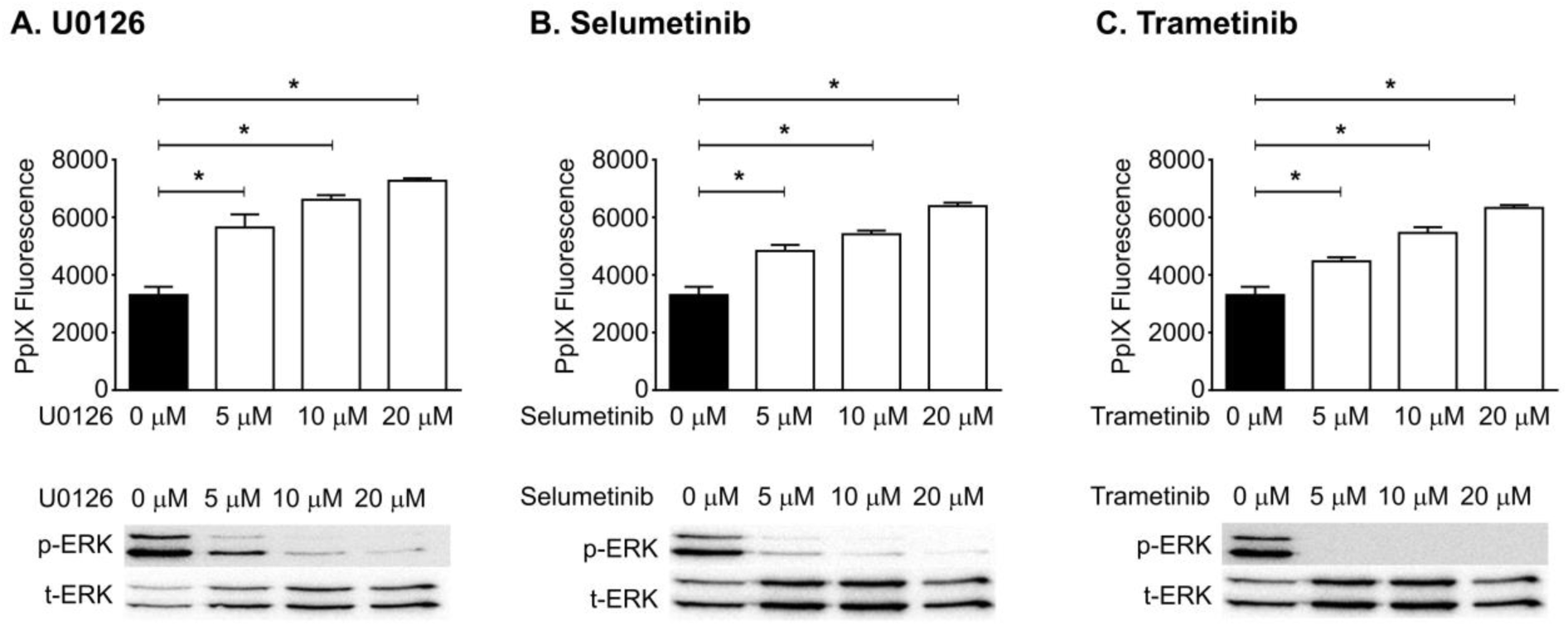
MEK inhibition increases PpIX accumulation in RasV12 cells. RAS V12 cells were treated with different concentrations of MEK inhibitors (A) U0126, (B) Selumetinib, or (C) Trametinib for 20 h, followed by 5 mM 5-ALA for 4 h. The plots show mean ± SD PpIX fluorescence from 3 independent experiments. *p<0.01 by one-way ANOVA with Turkey’s posthoc test. Western blots analysis of phosphorylated ERK (p-ERK) and total ERK (t-ERK) confirmed effective MEK inhibition.

To further determine the regulation of PpIX accumulation by mouse Ras/MEK *in vivo*, we used male HRAS transgenic mice, which spontaneously develop tumors in the thoracic region. This model was previously used to confirm that *in vivo* administration of 5-ALA results in tumor-specific accumulation of PpIX (19). Once the mice developed palpable tumors, they were treated i.p. with the MEK inhibitor U0126 or control vehicle; 5 h later the mice received i.p. 5-ALA, and tumor fluorescence was evaluated 2 h later (Fig. 3). PpIX fluorescence in tumor homogenates was significantly increased in tumors of mice treated with the MEK inhibitor and 5-ALA compared to those treated only with 5-ALA (Fig. 3A). This increase in PpIX fluorescence was evident in the tumors under blue light and confirmed by heat map analysis of the tumor images (Fig. 3B). These results indicate the presence of Ras/MEK-mediated regulation of PpIX accumulation in the *in vitro* and *in vivo* mouse systems.

**Fig 3.**
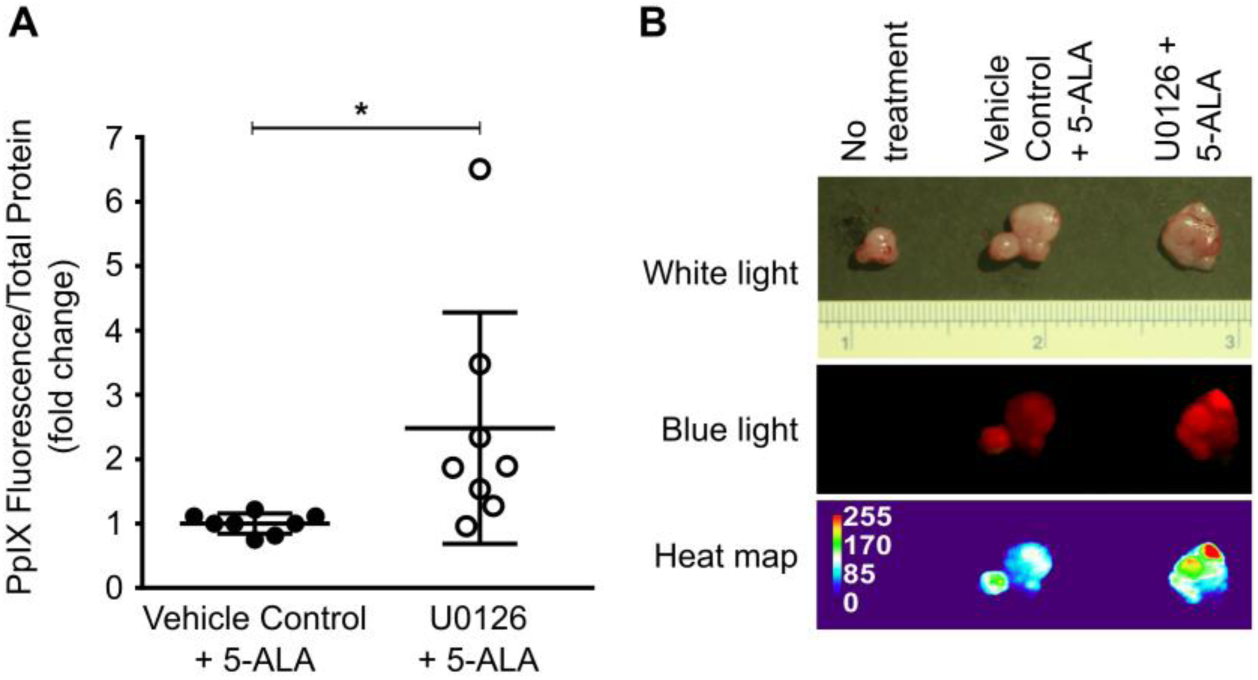
MEK inhibition enhanced tumor PpIX fluorescence in spontaneous tumors developed in HRAS transgenic mice. HRAS mice with palpable salivary gland tumors were treated i.v. with U0126 or control vehicle for 5 h and then i.p. with 5-ALA for 2 h. (A) Fold change in PpIX fluorescence in tumor homogenates normalized to total protein between mice treated with vehicle control +ALA and mice treated with U0126 + ALA is shown (n= 8/group). *p<0.05 by Student’s *t*-test. (B) Representative image of tumors in HRAS transgenic mice left untreated or treated with vehicle control + 5-ALA, or U0126 + 5-ALA and heat map generated based on tumor fluorescence.

### Ras/MEK regulates 5-ALA-induced PpIX accumulation via ABCB1 and FECH in the mouse fibroblast system

In our previous study, we demonstrated using human cancer cell lines, that Ras/MEK increases ABCB1 expression and FECH activity to promote PpIX efflux and its conversion to heme, respectively (14). Here, we determined whether these regulatory axes are present in the mouse fibroblast system (Fig. 4). Flow cytometry analysis showed that ABCB1 expression was higher in RasV12 cells compared to that in parental NIH3T3 cells (Fig. 4A). The expression of ABCB1 on RasV12 cells was significantly reduced (Fig. 4B) when they were treated with different concentrations of U0126 (5, 10, and 20 µM), suggesting that Ras/MEK regulates ABCB1 expression. To determine the involvement of the Ras/MEK-FECH axis, we evaluated FECH activity in RasV12 and NIH3T3 cells. FECH activity was significantly higher in RasV12 cells but was reduced to the level of the parental NIH3T3 cells when treated with a MEK inhibitor (Fig. 4C). Western blot analysis also demonstrated that MEK inhibition decreased FECH expression in RasV12 cells (Fig. 4D). These results confirm that the Ras/MEK-ABCB1 and Ras/MEK-FECH axes are involved in regulating 5-ALA-induced PpIX accumulation in the mouse fibroblast system.

**Fig 4.**
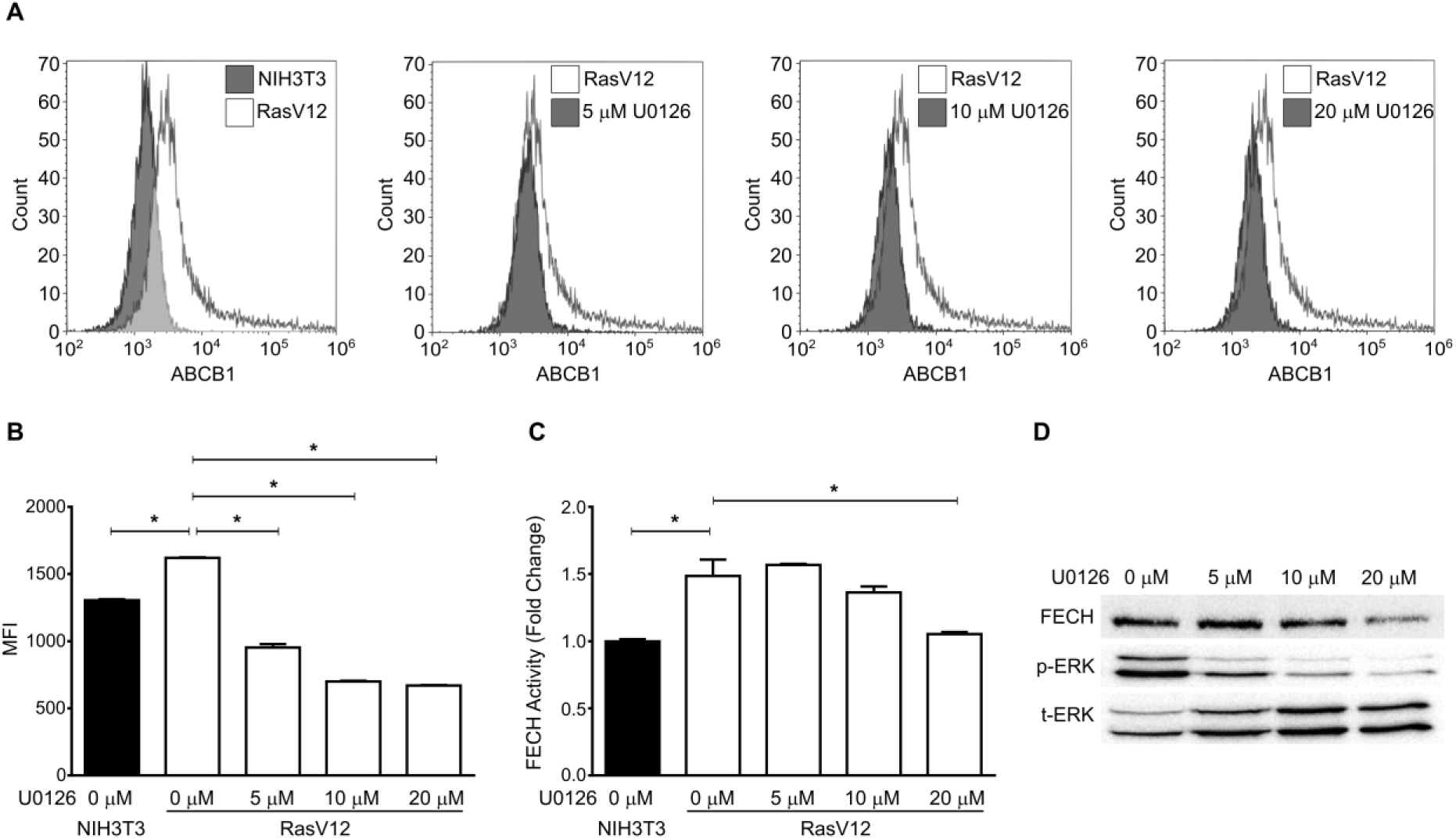
Oncogenic Ras/MEK regulates PpIX accumulation via ABCB1 and FECH. (A) Representative overlay histograms showing surface ABCB1 expression in NIH3T3 cells and RasV12 cells treated with or without different concentrations of MEK inhibitor, U0126. (B) Plot shows mean ± SD mean fluorescence intensity (MFI) from 3 independent experiments. (C) Mean ± SD relative FECH activity in NIH3T3 cells and RasV12 cells treated with or without different concentrations of U0126 from 3 independent experiments. (D) Representative western blot showing FECH expression and ERK phosphorylation levels in NIH3T3 and RasV12 cells treated with or without different concentrations of U0126. *p<0.01 by one-way ANOVA with Turkey’s posthoc test.

### RSKs are the downstream elements of Ras/MEK that regulate 5-ALA induced PpIX accumulation via ABCB1 expression

Activated Ras/MEK phosphorylates ERK1 and 2, which, in turn, activate MNKs, MSKs, and RSKs (18). As the interaction of RSK and ABCB1 was previously reported (20), we sought to determine whether RSKs are involved in the regulation of PpIX accumulation. First, we confirmed that MEK inhibition reduced RSK phosphorylation (p-RSK) in a dose-dependent manner, demonstrating that RSKs are the downstream elements of Ras/MEK in RasV12 cells (Fig. 5A). To determine whether RSKs regulate PpIX accumulation, we knocked down of RSK1, RSK2, RSK3, and RSK4, and evaluated PpIX accumulation. As shown by RT-PCR (RSK1, 3, and 4) and western blot analyses (RSK2), the RNAi knockdown efficiently reduced the expression of each RSK without affecting the expression of other RSKs (Fig. 5B). Knockdown of RSK2, RSK3, or RSK4, but not RSK1, promoted PpIX accumulation in RasV12 cells treated with 5-ALA, suggesting that these RSKs are the downstream elements of Ras/MEK responsible for decreasing PpIX accumulation (Fig. 5C). For further confirmation, we tested the effect of SL0101, a pan-RSK inhibitor (21), on PpIX accumulation in RasV12 cells (Fig. 5D). RSK inhibition significantly increased 5-ALA-induced PpIX accumulation. We also found that treatment with SL0101 reduced the expression of ABCB1 in RasV12 cells (Fig. 5E). These results demonstrate that Ras/MEK activates RSKs, which in turn up-regulate ABCB1 expression, resulting in increased PpIX efflux.

**Fig 5.**
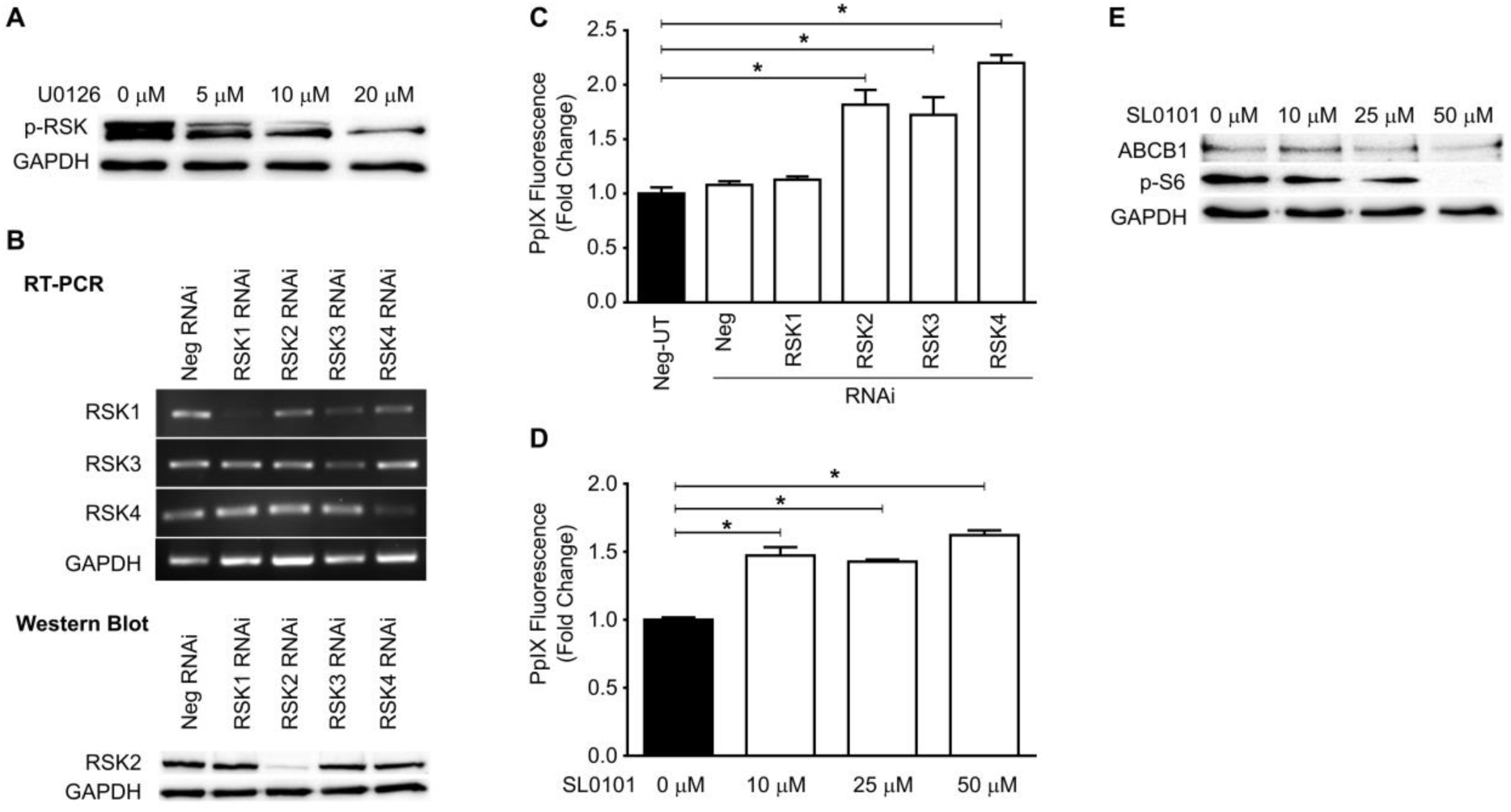
Oncogenic Ras regulates ABCB1 expression via RSKs. (A) Representative western blot showing RSK phosphorylation in RasV12 cells with or without MEK inhibition. (B) (Top) Representative gel image showing RSK1, RSK2, and RSK3 cDNA levels in RasV12 cells treated with or without various RSK siRNAs. (Bottom) Representative western blot showing RSK2 expression in RasV12 cells treated with or without various RSK siRNAs. (D, E) Fold change in PpIX accumulation in RasV12 cells (D) transfected with siRNA against RSKs and (E) treated with different concentrations of SL0101, a pan-RSK inhibitor. *p<0.01 by one-way ANOVA with Turkey’s posthoc test. (E) Representative western blots showing ABCB1 expression in RAS V12 cells treated with different concentrations of SL0101. p-s6 expression was used to confirm RSK inhibition, and GAPDH was used as the loading control.

### Ras/MEK-induced FECH activation is mediated through HIF-1α

Another downstream branch of the Ras/MEK-mediated regulation of PpIX accumulation is the conversion of PpIX to heme by FECH (14). A previous study indicated that FECH expression and activity are increased under hypoxia (22). It has also been shown that Ras/MEK increases the expression of HIF-1α, a key effector of cellular hypoxia (23). Therefore, we sought to determine the involvement of HIF-1α in the Ras/MEK-mediated regulation of PpIX accumulation. Consistent with previous reports (23, 24), MEK inhibition reduced the expression of HIF-1α in RasV12 cells treated with CoCl_2_, a known inducer of HIF-1α (Fig. 6A). When RasV12 cells were treated with HIF-1α inhibitor, FECH expression was reduced at the protein level but not the mRNA level (Fig. 6B), suggesting that HIF-1α regulates FECH expression at the post-transcriptional or translational levels. Furthermore, we found that FECH activity was significantly lower when RasV12 cells were treated with HIF-1α inhibitor (10 and 20 µM) (Fig. 6C). Finally, HIF-1α inhibition significantly promoted PpIX accumulation in RasV12 cells treated with 5-ALA in a dose-dependent manner (Fig. 6D). These results suggest that HIF-1α mediates Ras/MEK-mediated regulation of FECH, which increases the conversion rate of PpIX to heme.

**Fig 6.**
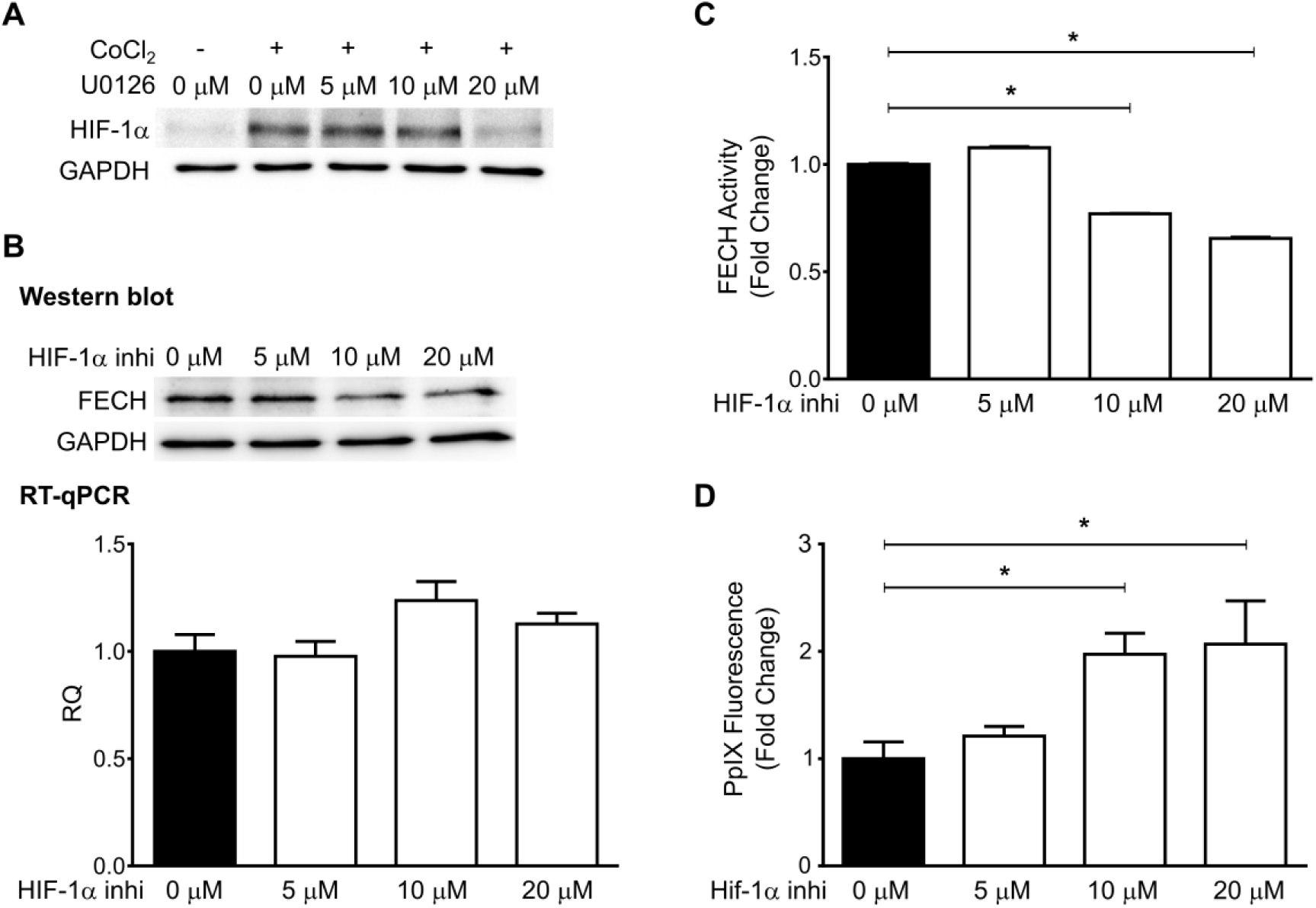
Oncogenic Ras regulates FECH activity via HIF-1α. (A) Representative western blot showing HIF-1α expression in RasV12 cells treated with or without CoCl_2_ and different concentrations of U0126. (B) Protein (Top) and mRNA (Bottom) levels of FECH in RasV12 cells treated with different concentrations of HIF-1α inhibitor. RQ: relative quantification. (C, D) Mean ± SD fold change in FECH activity (C) and PpIX accumulation (D) in RAS V12 cell lysates treated with different concentrations of HIF-1α inhibitor from 3 independent experiments. *p<0.01 by one-way ANOVA with Turkey’s posthoc test.

### Regulation of 5-ALA-induced PpIX accumulation via the Ras/MEK-RSK-ABCB1 and Ras/MEK-HIF-1α-FECH axes is present in human cancer cells

We identified that Ras/MEK regulates PpIX accumulation through two independent pathways – the RSK-ABCB1 and HIF-1α-FECH axes – in the mouse NIH3T3 fibroblast system. Next, we sought to determine whether these two pathways are involved in regulating 5-ALA-induced PpIX accumulation in human cancer cells. Four human cancer cell lines, DLD-1 (colon cancer), SNB-75 (glioblastoma), Hs 578T (breast cancer), and MDA MB 231 (breast cancer) were treated with an RSK inhibitor (SL0101), ABCB1 inhibitor (zosuquidar), or HIF-1α inhibitor (Fig. 7). RSK inhibition and HIF-1α inhibition enhanced 5-ALA-induced PpIX accumulation in all cell lines that were tested. Treatment with the ABCB1 inhibitor increased PpIX accumulation in all cell lines except MDA MB 231 cells. These results demonstrate that both Ras/MEK-RSK-ABCB1 and Ras/MEK-HIF-1α-FECH axes play critical roles in regulating PpIX accumulation in human cancer cells treated with 5-ALA.

**Fig 7.**
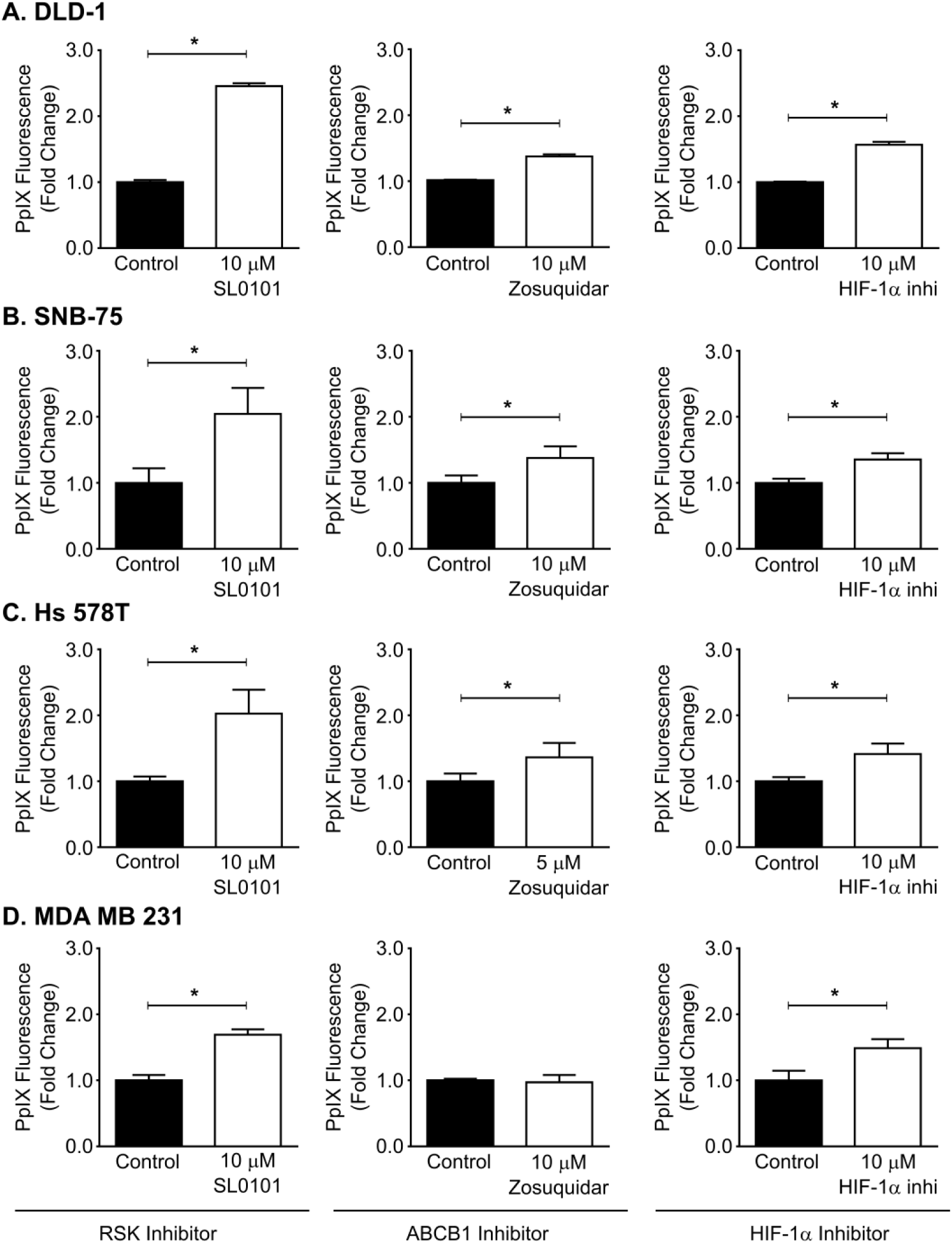
Inhibiting RSKs, ABCB1, and HIF-1α enhanced PpIX accumulation in human cancer cell lines. Human cancer cell lines DLD-1, SNB-75, Hs 578T, and MDA MB 231 were pre-treated with or without SL0101 (RSK inhibitor), Zosuquidar (ABCB1 inhibitor), or HIF 1α inhibitor for 20 h and then with 5-ALA for 4 h. Mean ± SEM fold change in PpIX fluorescence in cell lysate compared to controls is shown. *p<0.01 by one-way ANOVA with Turkey’s posthoc test.

## Discussion

The cancer-specific accumulation of PpIX is the key cellular mechanism used for the detection of tumors during surgeries (5-ALA-PDD) and for inducing cancer cell death by irradiating with light of specific wavelengths (5-ALA-PDT) (2–5). We previously demonstrated that inhibiting oncogenic Ras/MEK increases PpIX accumulation in cancer cells and promotes the efficacy of 5-ALA-PDD and PDT *in vitro* and *in vivo* (14, 17). We demonstrated that oncogenic Ras/MEK increases PpIX efflux and PpIX conversion to heme by promoting ABCB1 expression and FECH activity, respectively (14). However, it remained to be clarified how Ras/MEK regulates ABCB1 expression and FECH activity. To this end, we used mouse fibroblast cells transfected with RasV12 to systematically identify the downstream elements of Ras/MEK that regulate PpIX accumulation. Among the downstream pathways, we identified that RSKs are responsible for increasing ABCB1 expression and promoting PpIX efflux (Fig. 5 & 8). We also found that Ras/MEK activates FECH activity by increasing HIF-1α expression to promote the conversion of PpIX to heme (Fig. 6 & 8). Furthermore, we confirmed that the Ras/MEK-RSK-ABCB1 and Ras/MEK-HIF-1α-FECH axes are involved in the regulation of PpIX accumulation in human cancer cell lines (Fig. 7). Identifying the downstream elements is essential to further narrow down the possible molecular mechanisms of PpIX regulation by Ras/MEK. Further, the activation of RSKs and HIF-1α, in addition to Ras/MEK, may be useful as novel biomarkers in tumor biopsy samples to accurately predict the efficacy of 5-ALA-PDD and PDT in the clinical settings. Additionally, RSKs and HIF-1α could be developed as new therapeutic targets for combined treatment to promote the efficacy of 5-ALA-PDD and PDT.

**Fig 8.**
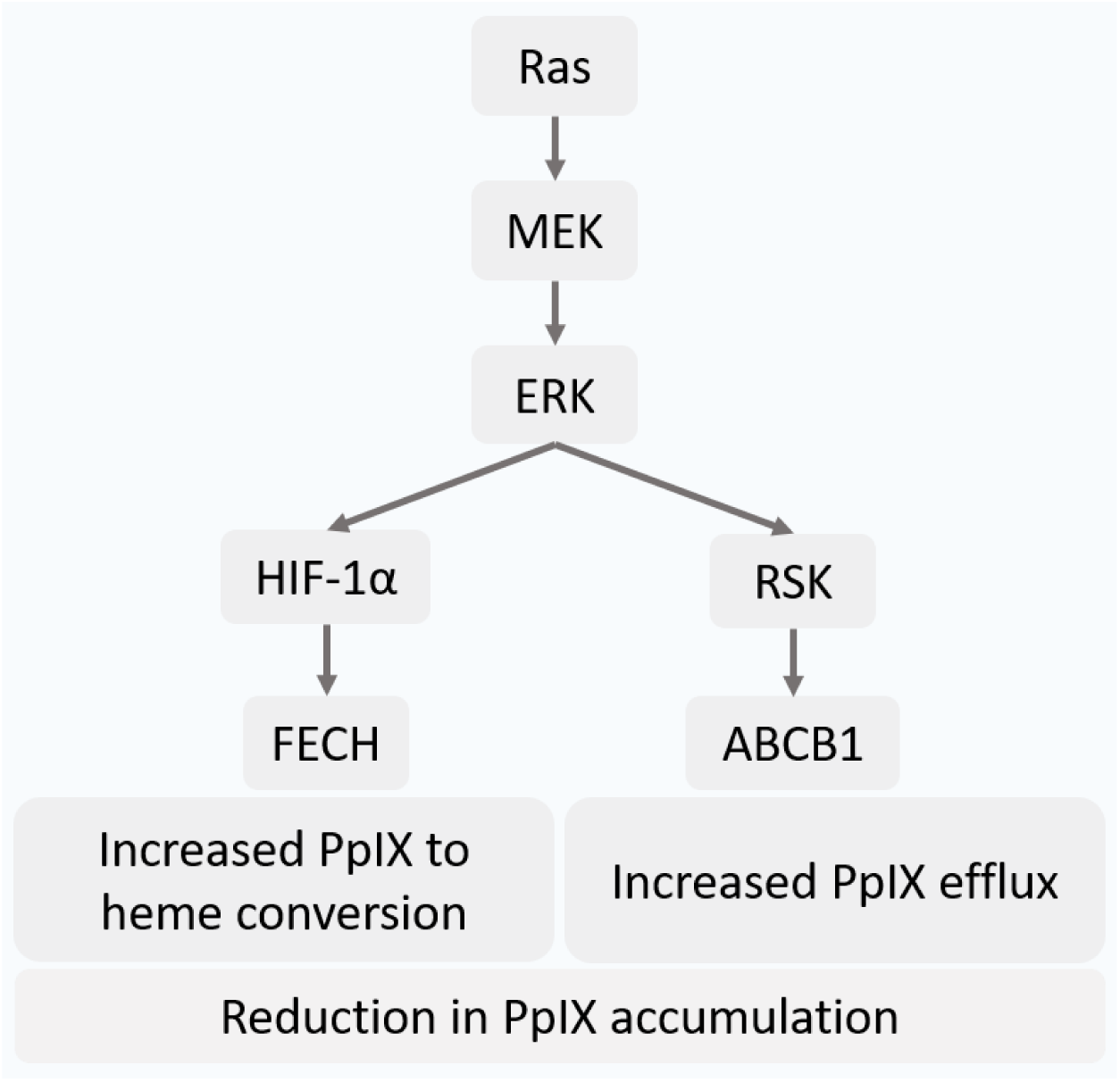
Reduction in PpIX accumulation through the Ras/MEK-HIF-1α-FECH and Ras/MEK-RSK-ABCB1 axes in cancer cells. Ras/MEK activation increases HIF-1α expression, which in turn increases the activity of FECH, the enzyme that catalyzes the conversion of PpIX to heme. Ras/MEK activation also upregulates ABCB1 expression through RSKs, which promotes the rate of PpIX efflux.

An important finding of this study is that treatment with a MEK inhibitor increased PpIX accumulation and tumor fluorescence in a mouse model of spontaneous cancer (Fig. 3). We previously used animal models with tumor implants of human or mouse cancer cell lines, which are highly homogenous, for evaluating PpIX fluorescence in tumors (14, 17). In contrast, tumors in the spontaneous mouse model consist of different stages of cancer cells and maintain tumor hierarchy, which closely resembles tumors in patients. Therefore, it was important to demonstrate that MEK inhibitor treatment could improve the efficacy of 5-ALA-PDD in the spontaneous mouse model.

RSKs (RSK1, RSK2, RSK3, and RSK4) are highly homologous isoforms but have distinct roles in regulating cellular processes, including growth, motility, survival, and proliferation (25). RSKs are also involved in transcriptional regulation by phosphorylating various transcription factors such as cyclic AMP response element-binding protein (CREB), c-Fos, c-Jun, and Serum response factor (Srf) (26–28). RSKs also stimulate cap-dependent translation by phosphorylating the translation initiation factor-4B (eIF4B) and the 40S ribosomal subunit protein S6 (rpS6) (29). In our RNAi experiments, we found that RSK2, RSK3, and RSK4, but not RSK1, are involved in regulating PpIX accumulation in Ras-transformed mouse fibroblast cells (Fig. 5). This is interesting as it is thought that RSK1 and RSK2 possess oncogenic functions that promote cancer cell growth, survival, and proliferation, while RSK3 and RSK4 are reported to have antitumor functions (30). This may be because RSKs are known to have unidentified, as well as overlapping functions (31). Furthermore, a previous study demonstrated that RSK1 increases ABCB1 expression by suppressing its ubiquitination and proteasomal degradation in human cancer cells (20), which is not consistent with our results. Discrepancies with the previous study might be due to the functional differences of RSKs between mice and humans. Therefore, a detailed analysis should be conducted to further clarify the roles of RSKs in ABCB1-mediated PpIX efflux.

Hypoxia is a common characteristic of advanced solid tumors, which reduces the efficacy of 5-ALA-PDD and PDT by different mechanisms (32). One of them is the increased expression of HIF-1α, which in turn reduces PpIX accumulation under hypoxic conditions (16, 33). HIF-1α is also a known transcriptional activator of the *FECH* gene (22). Furthermore, MEK activates the transcription of HIF-1α and induces resistance to cancer therapeutics under hypoxic conditions (23, 24). In this study, we demonstrated that oncogenic Ras/MEK reduces PpIX accumulation by activating the HIF-1α-FECH axis. As HIF-1α plays critical roles in developing resistance against other cancer therapeutics such as chemotherapy, radiotherapy, and immunotherapy, it is considered as a potential therapeutic target in cancer (34). There are several HIF-1 inhibitors currently being evaluated in phase II and III clinical trials (35). Therefore, once their safety and effectiveness are confirmed, it will be feasible to test the efficacy of combined 5-ALA-PDD and PDT with HIF-1α inhibitors in clinical settings. To this end, it is essential that further preclinical studies be conducted to determine the efficacy of the combined 5-ALA-PDD and PDT with HIF-1 inhibitors in animal models of cancer.

The activation of oncogenic signaling pathways is considered a major driving force for cancer-specific accumulation of PpIX (36). Ras pathways are one of the oncogenic signaling pathways that were previously reported to promote PpIX accumulation. Mouse fibroblast cells transformed with Ki-ras or v-raf showed higher PpIX accumulation than parental cells (9). Moreover, human breast epithelial cells transformed with Her2, an upstream element of Ras, also accumulated PpIX more efficiently than non-transformed cells (12). In contrast, we previously demonstrated that inhibition of oncogenic Ras/MEK increased PpIX accumulation in ∼60% of human cancer cell lines (14). In this study, we identified that the downstream elements of Ras/MEK, RSKs, and HIF-1α negatively regulate PpIX accumulation. These results agree with the previous study demonstrating that HIF-1α activation by ERKs contributes to 5-ALA-PDT resistance (16). Furthermore, similar roles of Ras/MEK have been observed in resistance to chemotherapy, where Ras/MEK activation increases drug-efflux functions of ABC transporters (37–39). As Ras signaling pathways regulate several other downstream elements including, phosphoinositide 3-kinases (PI3K), Ral guanine nucleotide exchange factors (RalGEF), and Ras-related C3 botulinum toxin substrate 1 (Rac1), it can be assumed that these pathways are involved in the promotion of PpIX accumulation by upregulating the enzymes of the heme biosynthesis pathway. In contrast, the Ras/MEK pathway reduces PpIX accumulation by activating PpIX efflux from the cells and its conversion to heme. The cancer-specific accumulation of PpIX is a well-established concept in the filed for decades, yet, the underlying cellular mechanisms are not completely understood. As elucidating the cellular mechanisms is key to improving the efficacy of 5-ALA-PDD and PDT, it is essential to further clarify the role of oncogenic transformation in cancer-specific PpIX accumulation.

## Experimental procedures

### Cells and reagents

NIH3T3 cells were obtained from the American Type Culture Collection (ATCC; Manassas, VA, USA). H-Ras-transformed NIH3T3 cells were generated in-house and was described previously (40, 41). Human colon cancer cell line DLD-1, glioblastoma cell line SNB-75, and breast cancer cell lines Hs 578T and MDA MB 231 were obtained from ATCC and were authenticated by STR DNA analysis (SickKids, Toronto). All cells were maintained in high glucose Dulbecco’s modified Eagle’s medium (DMEM) (Invitrogen, Ontario, Canada), supplemented with 10% fetal bovine serum (FBS) and antibiotic-antimycotic mixture (Invitrogen) (100 units/mL penicillin G sodium) at 37 °C and 5% CO_2_.

U0126 was purchased from Cell Signaling Technology (Danvers, MA); 5-Aminolevulinic acid from Sigma (Oakville, ON); HIF-1α inhibitor from Santa Cruz Biotechnology (Dallas, TX); ABCB1 inhibitor, zosuquidar from Selleckchem, and pan-RSK inhibitor, SL0101 from Calbiochem (Darmstadt, Germany). Anti-phospho-ERK-1/2 and anti-phospho RSK antibodies were purchased from Cell Signaling (Danvers, MA), anti-ABCB1 antibody from Alomone Labs (Israel), anti-FECH, anti-phospho-S6, anti-RSK2, anti-total ERK antibodies, and the FITC-tagged Anti-ABCB1 antibody (sc-55510) from Santa Cruz Biotechnology; anti-HIF-1α antibody (ab179483), and anti-GAPDH antibody were purchased from Abcam (US).

### Animal experiments

All animal care protocols were approved by the Institutional Animal Care Committee and were in accordance with the guidelines of the Canadian Council on Animal Care. B6; SJL-Tg(Wap-HRAS)69Lln Chr YSJL/J transgenic mice (hereafter referred to as HRAS mice) were obtained from Jackson Laboratory (JAX Mice Stock # 002409) and were housed in a barrier unit within the central animal care facility in the Health Sciences Center at Memorial University of Newfoundland. A unique feature of this mouse model is the incorporation of the HRAS (Harvey rat sarcoma viral oncogene homolog) oncogene on the Y chromosome, such that male mice expressing HRAS under the mammary tissue-specific whey acidic protein (Wap) promoter develop benign adenocarcinomas between 6-8 weeks of age (19, 42).

Male HRAS mice at approximately three months of age were used for the study. Once the mice developed palpable tumors, they were randomly assigned to one of the three groups – Control, ALA, or U0126+ALA (n=8 in each group). Mice in the U0126+ALA group were intraperitoneally (i.p) injected with U0126 (20 mg/kg body weight (BW) and those in control and ALA groups were administered i.p vehicle control (DMSO/saline). Five hours after U0126 treatment, mice in the ALA and U0126+ALA groups were administered i.p 5-ALA (200 mg/kg BW), and those in the control group were administered i.p saline. The mice were kept in their home cages and maintained in a darkened room for 2 h after ALA administration. The mice were then sacrificed by CO_2_ inhalation, and the tumors were excised. Tumor sizes varied from 2-10 mm in diameter and were distributed across the mammary tumor chain, primarily in the ventral, thoracic region.

The excised tumors were photographed using a Canon 6D camera fitted with a 35 mm lens mounted with a yellow 635 nm emission lens filter for fluorescence imaging. White light was used for bright field imaging, and blue light (405 nm) (Storz GmbH, Tuttlingen, Germany) was used for fluorescence imaging. Image analysis was carried out using ImageJ (NIH), and a heat map was generated using the HeatMap Histogram plugin (43). The tumors were homogenized in radioimmunoprecipitation assay (RIPA) buffer using a tissue homogenizer, and the homogenates were stored at −80 °C for further analysis.

### PpIX measurements

Cells (5 × 10^4^/well) plated in 24-well plates were treated with U0126, SL0101, HIF-1α inhibitor, zosuquidar, or DMSO (control vehicle) for 20 h, and then with 5-ALA for 4 h. The cells were lysed using RIPA buffer. PpIX fluorescence in cell and tumor lysates was measured using a Synergy Mx Fluorescence plate reader (BioTek Instruments Inc. VT) with a 405 nm excitation/630 nm emission filter. The total protein in the tumor lysate was determined using the BCA total protein assay kit (Thermo Scientific) following the manufacturer’s instructions, and the PpIX fluorescence was normalized to total protein.

### Protein expression analysis

Cells treated with or without the inhibitors were lysed using RIPA buffer supplemented with aprotinin (Sigma), and Halt™ Protease Inhibitor Cocktail (100X) (Thermo Scientific). The protein samples were subjected to sodium dodecyl sulfate-polyacrylamide gel electrophoresis (PAGE) and transferred to a nitrocellulose membrane (Bio-Rad, Canada). The expression levels of ABCB1, FECH, p-ERK, t-ERK, RSK2, p-RSK, p-S6, or HIF-1α were determined using the corresponding antibodies as described previously (44). For determining the cell surface expression of ABCB1, cells (10^7^ cells) treated with or without different concentrations of U0126 were suspended in 100 µl flow cytometry buffer (PBS containing 0.5% bovine serum albumin (BSA) and 2 mM EDTA), and incubated with 10 µl anti-ABCB1-FITC antibody in the dark for 15 min at 4 °C. The cells were then washed with and re-suspended in 500 µl flow cytometry buffer and analyzed using a BC CytoFLEX flow cytometer (Beckman Coulter).

### Knockdown of RSK using siRNA

The negative control siRNA and siRNA against human RSK1, RSK2, RSK3, and RSK4 were purchased from Santa Cruz Biotechnology. A day before transfection, RasV12 cells (2.5 × 10^4^ cells/well) were plated in 24-well plates. The cells were transfected with 10 pmol siRNA using Lipofectamine RNAi MAX (Life Technologies) in serum-free medium. The transfection was repeated 24 h later. Protein and RNA samples were collected 48 h after the second transfection. RSK2 protein expression was determined by western blot analysis, and the expression levels of RSK1, RSK3, and RSK4 were determined by RT-PCR.

### RT-PCR and RT-qPCR

Total RNA was isolated from cells using TRIzol (Invitrogen) according to the manufacturer’s instructions. RNA (0.5 µg) was reverse transcribed (RT) to cDNA from random hexamers using the ReverAid H Minus First Strand cDNA Synthesis Kit (Thermo Scientific). Semi-quantitative RT-PCR for RSK elements was performed on the cDNA using the following primers – RSK1: (forward) 5′-GAGAGACATCCTCGCTGACG-3′, (reverse) 5′-TGCCTAGCTTCGCCTTCAAA-3′; RSK3: (forward) 5′-CTCCCAAGGGG TTGTCCATC-3′, (reverse) 5′-CACGGGTGCTTCAACACTTG-3′; RSK4: (forward) 5′-GTTGGCTGGCTACACTCCAT-3′, (reverse) 5′-ATATGGTGCTGCCACTGCTT-3′; GAPDH: (forward) 5′-GGGTGGAGCCAAACGGGTCA-3′, (reverse) 5′-GGAGTTGCTGTTGAAGTCGCA-3′. Cycling conditions were: 94 °C for 5 min followed by 34 cycles of 94 °C for 1 min, 66 °C for 1 min and 72 °C for 1 min; followed by a final extension of 10 min at 72 °C. PCR products were detected by agarose gel electrophoresis using RedSafe™ Nucleic Acid Staining Solution (iNtRON Biotechnology, Inc.).

Quantitative RT-PCR (RT-qPCR) for FECH was performed in triplicate on the cDNA using the following primer set. FECH: (forward) 5′-GATGGAGAGAGATGGACTAGA-3′, (reverse) 3′-TGCCTGCTTCACCACCTTCTT-5′; GAPDH: (forward) 5′-GATGGAGAGAGATGGACTAGA-3′, (reverse) 3′-TGCAAAGCACTGGATGAG-5′. Primers were validated using a 5-point, 5-fold dilution series. The absence of non-specific amplification was confirmed by observing a single peak in the melt-curve analysis, confirmation of the expected amplicon size by agarose gel analysis, and the absence of amplification in the no template control. qPCR was then performed in triplicate on the StepOne Plus (Applied Biosystems CA, USA) using the *power*SYBR^®^ Green PCR Master Mix (Life Technologies LTD, UK). Cycling conditions were: 50 °C for 3 min, 95 °C for 5 min followed by 40 cycles of 95 °C for 15 s, 60 °C for 30 s, and 40 °C for 1 min. The relative FECH mRNA level was calculated using the 2^-ΔΔCT^ method.

### FECH activity assay

FECH activity was measured by aerobic enzymatic formation of zinc-protoporphyrin IX (Zn-PpIX) using a modified van Hillegersberg method as described previously (45, 46). Briefly, cell lysates were incubated with 200 µM PpIX (Sigma Aldrich) in 200 µL assay buffer (0.1 M Tris-HCl, 1 mM palmitic acid (Sigma Aldrich) and 0.3% v/v Tween 20, pH 8.0) and then 50 µL 2 mM zinc acetate solution was added. The mixture was incubated at 37 °C for 60 min. The reaction was terminated by adding 500 µL ice-cold stop buffer (1 mM ethylenediaminetetraacetic acid (EDTA) in 30:70 DMSO/ methanol). The reaction mixture was centrifuged at 14,000 *×g* for 10 min, and Zn-PpIX in the supernatant was measured using a Synergy Mx Fluorescence plate reader (BioTek Instruments Inc. VT) with a 405 nm excitation/590 nm emission filter. Heat-inactivated cell lysates were included as a negative control.

### Statistical analysis

Statistical analyses were performed using Prism 7.0 (GraphPad). Student’s *t*-test was used for inter-group comparison, and one-way ANOVA with Tukey’s posthoc test was used to compare between multiple groups. p < 0.01 and p < 0.05 were considered statistically significant for *in vitro* and *in vivo* experiments, respectively.

## Data availability

All data are contained within the manuscript.

